# Rhizospheric bacteria from the Atacama Desert hyper-arid core: cultured community dynamics and plant growth promotion

**DOI:** 10.1101/2024.01.04.574204

**Authors:** Juan Castro-Severyn, Jonathan Fortt, Mariela Sierralta, Paola Alegria, Gabriel Donoso, Alessandra Choque, Marcela Avellaneda, Coral Pardo-Esté, Claudia P. Saavedra, Alexandra Stoll, Francisco Remonsellez

**Affiliations:** Laboratorio de Microbiología Aplicada y Extremófilos, Facultad de Ingeniería y Ciencias Geológicas, Universidad Católica del Norte, Antofagasta, Chile; Centro de Investigación Tecnológica del Agua y Sustentabilidad en el Desierto-CEITSAZA, Universidad Católica del Norte, Antofagasta, Chile; Laboratorio de Ecología Molecular y Microbiología Aplicada, Departamento de Ciencias Farmacéuticas, Facultad de Ciencias, Universidad Católica del Norte, Antofagasta, Chile; Laboratorio de Microbiología Molecular, Facultad de Ciencias de la Vida, Universidad Andres Bello, Santiago Chile; Centro de Estudios Avanzados en Zonas Áridas (CEAZA), La Serena, Chile

**Keywords:** Atacama Desert, *D. spicata*, *S. foliosa*, microbial cultures, rhizobiome

## Abstract

The Atacama Desert is the oldest and driest desert on Earth, with environmental conditions including great temperature variations, high UV-radiation, drought, high salinity, making it a natural laboratory to study the limits of life and resistance strategies. However, it shows great biodiversity harboring vast forms of adapted life and can be used as a model of desertification processes. While desertification is increasing as result of climate change and human activities, is necessary to optimize soil and water usage, where stress-resistant crops are possible solutions. As many studies have revealed the great impact of rhizobiome over plant growth efficiency and resistance to abiotic stress, we set up to explore the rhizospheric soils of *Suaeda foliosa* and *Distichlis spicata* from the Atacama Desert. By culturing these soils and using 16S rRNA amplicon sequencing, we address the community taxonomy composition dynamics, the stability through time and the ability to promote lettuce plants growth. The rhizospheric soil communities were dominated by the families Pseudomonadaceae, Bacillaceae and Planococcaceae for *S. foliosa* and Porphyromonadaceae and Haloferacaceae for *D. spicata*. Nonetheless, the cultures were completely dominated by the Enterobacteriaceae family (up to 98%). Effectively, lettuce plants supplemented with the cultures showed greater size and biomass accumulation, we identify 12 candidates that could be responsible of these outcomes, of which 5 (*Enterococcus, Pseudomonas, Klebsiella, Paenisporosarcina* and *Ammoniphilus*) were part of the built co-occurrence network, being *Klebsiella* a major participant. We aim to contribute to the efforts to characterize the microbial communities as key for the plant’s survival in extreme environments, and as a possible source of consortia with plant growth promotion traits aiming agricultural applications.

**IMPORTANCE:** The current scenario of climate change and desertification represents a series of incoming challenges for all living organisms, also as the human population grows rapidly, so is rising the demand for food and natural resources; thus, it is necessary to make agriculture more efficient by optimizing soil and water usages thus ensuring future food supplies. Particularly, the Atacama Desert (northern Chile) is considered the most arid place on Earth as a consequence of geological and climatic characteristics, such as the naturally low precipitation patterns and high temperatures, which makes it an ideal place to carry out research that seeks to aid agriculture to the future sceneries, which are predicted to resemble these. The use of microorganism consortia from plants thriving under these extreme conditions to promote plant growth, improve crops and make "unsuitable" soils farmable is our main interest.

**TWEET:** Cultures of rhizospheric soils from Atacama Desert resilient plants were enriched in *Klebsiella*, *Bacillus* and *Brevibacillus* which promoted lettuce growth

## INTRODUCTION

The Atacama Desert is in the dry subtropical climate belt between 18 °S and 27 °S, extending from the coastal edge to the Andean Mountain complex, and is the oldest and driest desert on Earth (Berger and Cooke, 1997; Clarke, 2006). This ecosystem is considered a natural laboratory, due its environmental co-occurring conditions, including high variations in temperature, UV radiation, hydric stress, presence of metal(oids), among many others (Navarro-González et al., 2003). Also, soil weathering, leaching and water erosion rates are slow in the area (Ewing et al., 2006, Ewing et al., 2008). Despite hostile conditions, the Atacama Desert harbors a vast adapted life forms making it possible to find great biodiversity. Among these, hypolithic cyanobacteria (Warren-Rhodes et al., 2006), non-lichenized fungi (Conley et al., 2006), lichens (Rundel, 1978), cacti (Rundel et al., 1991), and even shrubs and trees (Fletcher et al., 2012) have been reported. Moreover, the Atacama Desert hyper-arid core is a hotspot for studies on astrobiology and poly extremophile life (Hock et al., 2007; Cabrol et al., 2009; Parro et al., 2011). This ecosystem is under constant hydric stress, as the annual mean precipitation is lower than 2 mm, and there are years that do not receive any rain (Fuentes et al., 2022). This makes this area an ideal model for understanding the drought resistance bases displayed by these organisms.

Highly adapted microbial taxa thrive in these hyper-arid environments by having multiple adaptations for effective colonization and stress tolerance (Tian et al., 2017; Torres-Cortés et al., 2018). Particularly, members of the Firmicutes, Bacteroidota and Actinobacteria phyla (Rubrobacterales, Actinomycetales, and Acidimicrobiales) have been associated with halite nodules and soil samples from these environments as they can develop with low humidity, high soil salinity, and high solar radiation conditions (Crits-Christoph et al., 2015, 2016; Piubeli et al., 2015; Fuentes et al., 2022). Moreover, in this type of extreme environment, the presence of plants increases the organic matter levels (Vinton and Burke, 1995; Burke et al., 1999), where symbiotic interactions are established between microorganisms and plant roots, thus maintaining the nutrient cycling in soils and optimize resources (Conley et al., 2006; Martirosyan et al., 2016; Lopez and Bacilio, 2020; Jones et al., 2023). Also, plants form patches of vegetation that generate spatial heterogeneity and changes in pH at different scales in the soil due to moisture and nutrient retention (Yin et al., 2010; Wang et al., 2019; Wang et al., 2019), which also generates changes in pH. These symbiotic relationships are further promoted in extreme environments as they can increase the species survival under stress conditions, independently of their innate characteristics (Puente et al., 2009; Trivedi et al., 2020). Some microorganism traits from which plants can benefit from are salt tolerance, zinc potassium and phosphorus solubilization, ammonia siderophores, phytohormones and secondary metabolites production (Lopez-Lozano et al., 2020). Nonetheless, the key groups and the role they would be playing in the interaction under these particular extreme environment remains unclear.

The use of microorganisms that contribute with beneficial traits to plant development on increased yield can be a successful strategy to contribute to the current agricultural industry crisis (Verma et al., 2017; Odoh et al., 2020; Rizvi et al., 2021). Moreover, strains isolated from plant species living in highly challenging and stressful environments have been evaluated for their protective capacity against stress conditions or as growth promoters in crops (Inostroza et al., 2017; Maza et al., 2019), for instance microorganisms can induce drought resistance, improve the photosynthetic rates, promote phytohormones, and increase biomass (Glick et al., 2007; Bal et al., 2013; Giauque and Hawkes, 2013; Torres-Diaz et al., 2016), including agricultural relevant crops such as *Lactuca sativa, Hordeum vulgare, Oryza sativa* and *Chenopodium quinoa* (Waller et al., 2005; Redman et al., 2011; Molina-Montenegro et al., 2016; González-Teuber et al., 2018; Santander et al., 2020). Despite the fact that most of these investigations are based on isolated bacteria or yeasts strains, there is compelling evidence for the use of microbial consortia as biofertilizer, promoting growth in challenging conditions, increasing crop yield, nutrient uptake, salt stress (Zayadan et al., 2014; Dal Cortivo et al., 2018; Odoh et al., 2020; Redondo-Gómez et al. 2021; Seenivasagan et al., 2021; Fortt et al., 2022). Moreover, the effects of biofertilizers on soil and rhizosphere are still under-characterized and its significance on ecological functions (Sharma et al., 2012). Despite this, the monitoring of direct soil cultures throughout time has not been addressed until now, as well as the characterization of the substrate microbial community composition post biofertilization experiment.

Our group identified an oasis in the Yungay area of the Atacama Desert Hyper-arid Core (The Aguas Blancas Basin), this place is uncoupled from the coastal fog due to a mountain range and rainfall is currently insufficient to support vascular plants in most of the area (Rech et al., 2003). Nonetheless, a small oasis or fertile island can be found, harboring high density of plants and shrubs that thrive facing the high salinity soils and extreme drought (Fletcher et al., 2012). The most abundant plant is *D. spicata,* which is very adapted to grow in alkaline saline soils and also contributes to the construction of mounds around its individuals, favoring localized elimination of salts by capillary action and evaporation. While *Suaeda foliosa* is a perennial decumbent plant that does not require large amounts of water for development since it is specialized in capture and retaining it (Conticello et al., 2002; Pelliza et al., 2005; Pfeiffer et al., 2018). In this study we set up to characterize the microbial community composition of *S. foliosa* and *D. spicata* rhizospheric soils, then culture these soils to monitoring the dynamic and stability through time and several subcultures to finally test their ability to promote *L. sativa* growth and determining which taxa could be playing a key role in the lettuce improvement.

## MATERIALS AND METHODS

### Field trip and sample collection

A sampling expedition was carried out during September 2022 to the Yungay Hyper-Arid Core area, located in the Antofagasta Region of Chile (between approximately 22°S and 26°S; Fuentes et al., 2022; Jones et al., 2023). The mean annual rainfall for this area is around 2.0 mm (making it the driest zone in the desert) and long-term climate data indicate that it is also the driest non-polar desert on Earth, due low rainfall, lack of water, high temperatures and evapotranspiration rates as well as the prevalent cloudless condition, low total ozone column, the highest surface ultraviolet (UV) radiation and total solar irradiance recorded in the planet (Navarro-González et al., 2003; 2013; Azua-Bustos et al., 2017; Ritter et al., 2018). Despite all these conditions we located a small oasis area or fertile island (24°3’29.93"S and 69°49’33.25"O; Figure 1: upper panel) with the presence of three plants species: *Distichlis spicata* which is a grass (the most abundant one) also known as "desert saltgrass"; *Suaeda foliosa* is a bush (representing much lower coverage) and finally one specimen of *Prosopis tamarugo* which is a leguminous deciduous tree. Rhizospheric soil samples were taken for *D. spicata* and *S. foliosa* (3 for each) in an aleatory sampling, trying to cover as much of the area as possible, we dug with a (70% ethanol sterilized) hand shovel, the closest to the plant root possible and approximately 10 cm deep and the soil samples were collected in 50 ml sterile falcon tubes and immediately transported to the laboratory.

**Figure 1.**
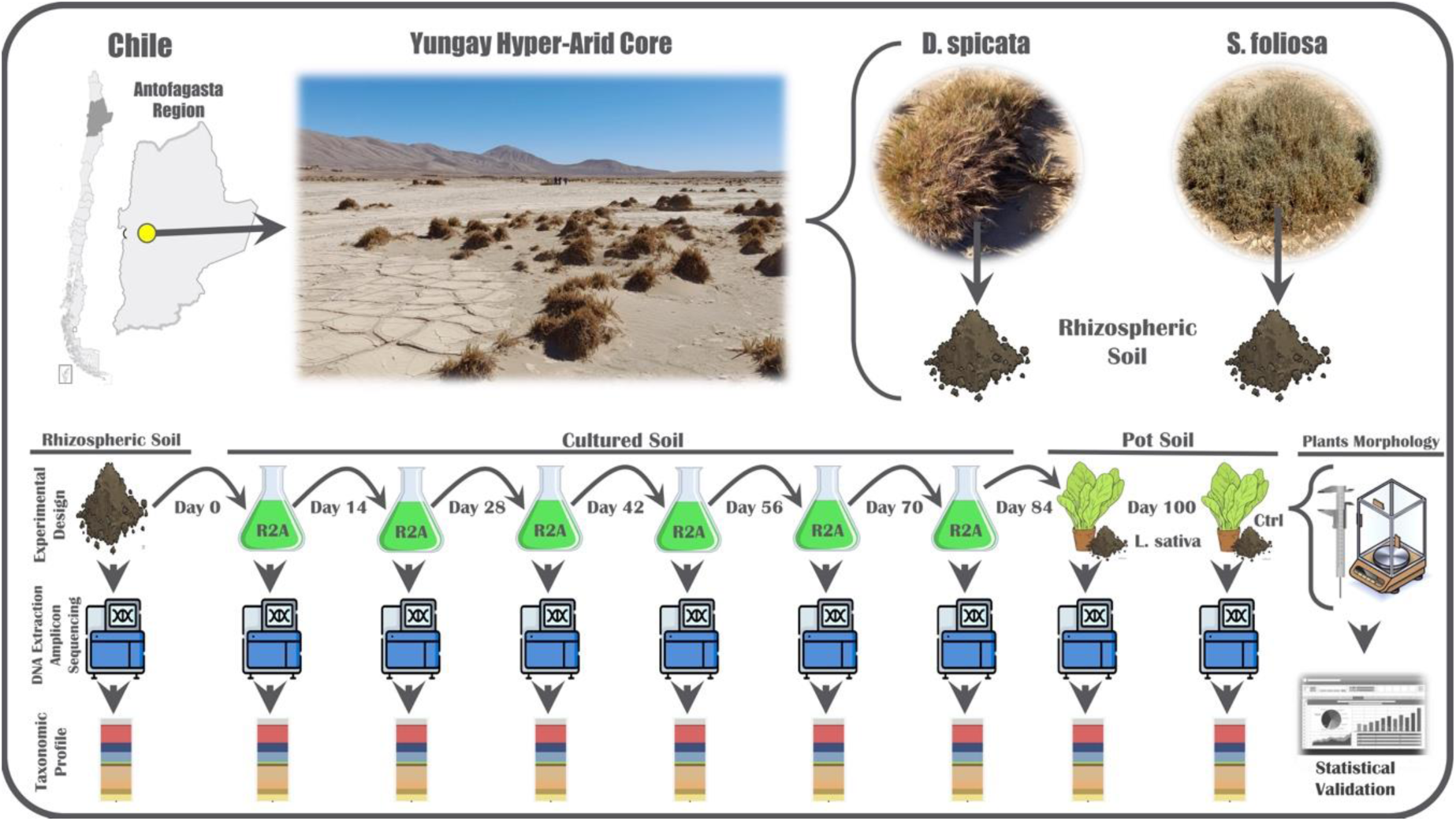
Sampling and experimental design scheme. In the upper panel we can see the location of Yungay Hyper-Arid Core (22°S and 26°S) and the fertile island (24°3’29.93"S and 69°49’33.25"O) and the sampled plant species photos. In the lower panel we can see the experimental design following the rhizospheric soil subcultures through time and the points in which DNA was extracted to determine the taxonomic composition.

### Soil Cultures

Approximately 0.5 gr of rhizospheric soil were inoculated in flasks with 20 ml of sterile Reasoner’s 2A (R2A) medium (Reasoner and Geldreich, 1985: 0.5 g/l yeast extract, 0.5 g/l proteose peptone, 0.5 g/l casamino acids, 0.5 g/l dextrose, 0.5 g/l soluble starch, 0.5 g/l sodium pyruvate; 0.3 g/l dipotassium phosphate and 0.03 g/l magnesium sulphate, pH 7.2) and cultured at 20°C with constant agitation (120 rpm). After 14 days, 1 ml of the grown culture was inoculated in a new flask containing 20 ml of fresh sterile R2A medium to continue culturing for 14 more days under the same conditions (Figure 1: lower panel); the remaining culture was centrifuged (3,000 g, 10 min) to discard the supernatant and the pellets were stored at −80°C until DNA extraction. This process was repeated four more times until reaching day 84, day on which the cultures were divided in three parts, the first part was pelleted and stored at −80°C until DNA extraction. The cultures growth was monitored weekly through the OD_600_ reading.

### Potted Plants Experiment

For this experiment, 15 days old green leaf lettuce (*Lactuca sativa* var. crispa) seedlings were obtained from Vivero La Portada (Antofagasta, Chile). The seedling roots were carefully washed and inoculated with a culture suspension for 1 h (the suspension was prepared with by pooling the second parts of the day 84 cultures and dilute at a 1:10 ratio with sterile distilled water). Then, inoculated and control (non-inoculated) seedlings were transplanted to 250 ml pots with a sterilized substrate mixture (2:1:1, leaf mold/sand/perlite) and grown for 16 days in an open-air shader with a natural photoperiod of 12 h/12 h approx. Four treatments were tested: *L. sativa* plants inoculated with *D. spicata* culture; *L. sativa* plants inoculated with *S. foliosa* culture; *L. sativa* plants without any inoculation (Control) and *L. sativa* plants supplemented with diluted R2A sterile media (ControlMed). Each treatment consisted in 12 independent and randomized replicates (plants). Regular irrigation was carried out every 48 h with 50 ml of distilled water at the afternoon. At day 8 after transplant, a second inoculation with 50ml of culture suspension (prepared by pooling the third parts of the day 84 cultures) was carried out, to the corresponding pots. After, normal irrigation with water continued in the same way until day 100, when the lettuce plants were harvested and corresponding soil samples were taken from selected pots and stored at −80°C until DNA extraction.

### Plants Morphological Evaluation

After the harvest, plant roots were carefully washed with distilled water to remove any remaining substrate and then they were air-dried over paper towels for 30 min. Later, plants were weighted (fresh weight), the longest root and the second “true leaf” lengths were measured for each plant. Next, each plant was deposited inside a paper bag and these were dried in a laboratory stove at 70°C for 48 hours and then weigh them again (dry weight) to calculate the dry matter content. All data were carefully recorded and the statistical significance was tested through one-way ANOVA with post hoc Tukey HSD for all comparisons (GraphPad 5.0: Prism) and visualizations were made using R package ggplot2 (Wickham, 2016).

### DNA Extraction and Amplicon Sequencing

Total DNA was extracted from *S. foliosa* and *D. spicata* rhizospheric soil, culture pellets and potted plant experiment soils using the E.Z.N.A. Soil DNA Extraction Kit (Omega Bio-tek, USA) according to the manufacturer’s instructions. DNA integrity, quality, and quantity were verified by 1% agarose gel electrophoresis, OD_260/280_ ratio spectroscopy, and fluorescence using a Qubit 3.0 fluorometer along with the Qubit dsDNA HS assay kit (Thermo Fisher Scientific, USA). Next, DNA samples were sent to the Environmental Sample Preparation and Sequencing Facility at the Argonne National Laboratory (Illinois - USA) for amplification of the bacterial 16S rRNA gene V4 region (∼250 bp) using the 515F and 806R primers (Caporaso et al., 2011), construction of 151 bp paired-end libraries and sequencing on a MiSeq (Illumina) platform.

### Taxonomic composition and diversity analysis

This analysis was conducted in R v4.2.2 and RStudio v1.3.1093 following the DADA2 v1.26.0 R package pipeline (Callahan et al., 2016), in order to infer amplicon sequence variants (ASVs) present in each sample. Briefly, the reads were evaluated for quality control and subsequently trimmed (Ns = 0, length ≥ 130bp, expected errors ≤ 2), followed by dereplication, denoising and merging of paired reads. Subsequently, the ASVs (amplicon sequence variants) table was built with 97% clustering, the chimeras were removed, and taxonomic assignment was carried out against the Silva v138 (Quast et al., 2012) database with the Ribosomal Database Project’s (RDP) naive Bayesian classifier (Wang et al., 2007). ASVs identified as Eukarya, Chloroplast, and Mitochondria were removed. Moreover, a multi-sequence alignment was created with DECIPHER v2.26.0 (Wright, 2016), to infer phylogeny using FastTree v2.1.11 (Price et al., 2009). Furthermore, a phyloseq-object (containing the ASVs, taxonomy assignment, phylogenetic tree, and the samples meta-data) was created using the R package Phyloseq v1.42.0 (McMurdie and Holmes, 2013). Replicates per condition were averaged by day or stage. The ASV counts were normalized by variance-stabilizing transformation using the R package DESeq2 v1.38.3 (Love et al., 2014). Alpha diversity indices were calculated using the Microbiome v1.20.0 and Btools v0.0.1 packages. Plots were generated using the ggpubr v0.6.0 package with comparisons between plant species using the Wilcoxon test (P < 0.05) and the statistical significance of the communities variation throughout the experiments was evaluated with ANOVA and Kruskal Wallis test. Taxonomy composition and relative abundance plots were generated using the ggplot2 v3.4.1, fantaxtic v0.2.0 and ampvis2 v2.7.35 (Andersen et al., 2018) R packages. Moreover, candidate taxa were identified by filtering with the following criteria to keep those that: 1) were detected in the rhizospheric soils of Yungay; 2) were detected (maintained) throughout the 84 days cultures; 3) were detected in the pots soil (after the bio-fertilization experiments of *L. sativa*) and 4) had a higher abundance in the bio-fertilized pots soil, regarding to the control pots soil. Finally, co-occurrence networks were constructed by agglomerating the phyloseq object at best hit using the microbiomeutilities v1.00.11 R package (Lahti et al., 2017), the network was estimated using the SpiecEasi v0.1.4 (Kurtz et al., 2015) R package (neighborhood selection model) and visualized with GGally v1.5.0 (Schloerke et al., 2018) R package.

### Data availability

The whole amplicon sequencing raw data sets have been deposited at DDBJ/ENA/GenBank under the Bioproject: PRJNA971922.

## RESULTS

The direct inoculation of rhizospheric soil from Yungay into culture medium effectively resulted in the propagation of microorganisms. This microbial growth was evident with increased medium turbidity, and although there was growth in all the subcultures, the DO_600_ monitoring every 14 days revealed a negative slope when comparing the cultures of both plant species throughout time (Supplementary Figure S1). Initially, at day 14 the cultures from both plants displayed very different values (being higher those from *D. spicata*) this difference progressively decreased as the growth rate of both cultures become slower and from day 56, they become equivalent.

Alpha diversity metrics were monitored throughout the experiment and compared between the stages and as expected, all evaluated indices decrease throughout the experiment (Figure 2). No significant differences were observed between both plants in the three used metrics, even when considering only the rhizospheric soil communities of *S. foliosa* and *D. spicata* (Supplementary Figure S2). Furthermore, changes among the stages or through time are evident in the three indices. The Shannon diversity index decreased progressively in cultures of microorganisms obtained from the Yungay rhizospheric soils towards the day 56 culture, subsequently they remained roughly stable until day 84, reaching the highest value on the lettuce pot soils. Moreover, the same pattern is observed for the Chao1 index and the phylogenetic diversity but to a lesser extent. In addition, the soil from the control lettuce pots displayed similar values for the three metrics to the inoculated ones.

**Figure 2.**
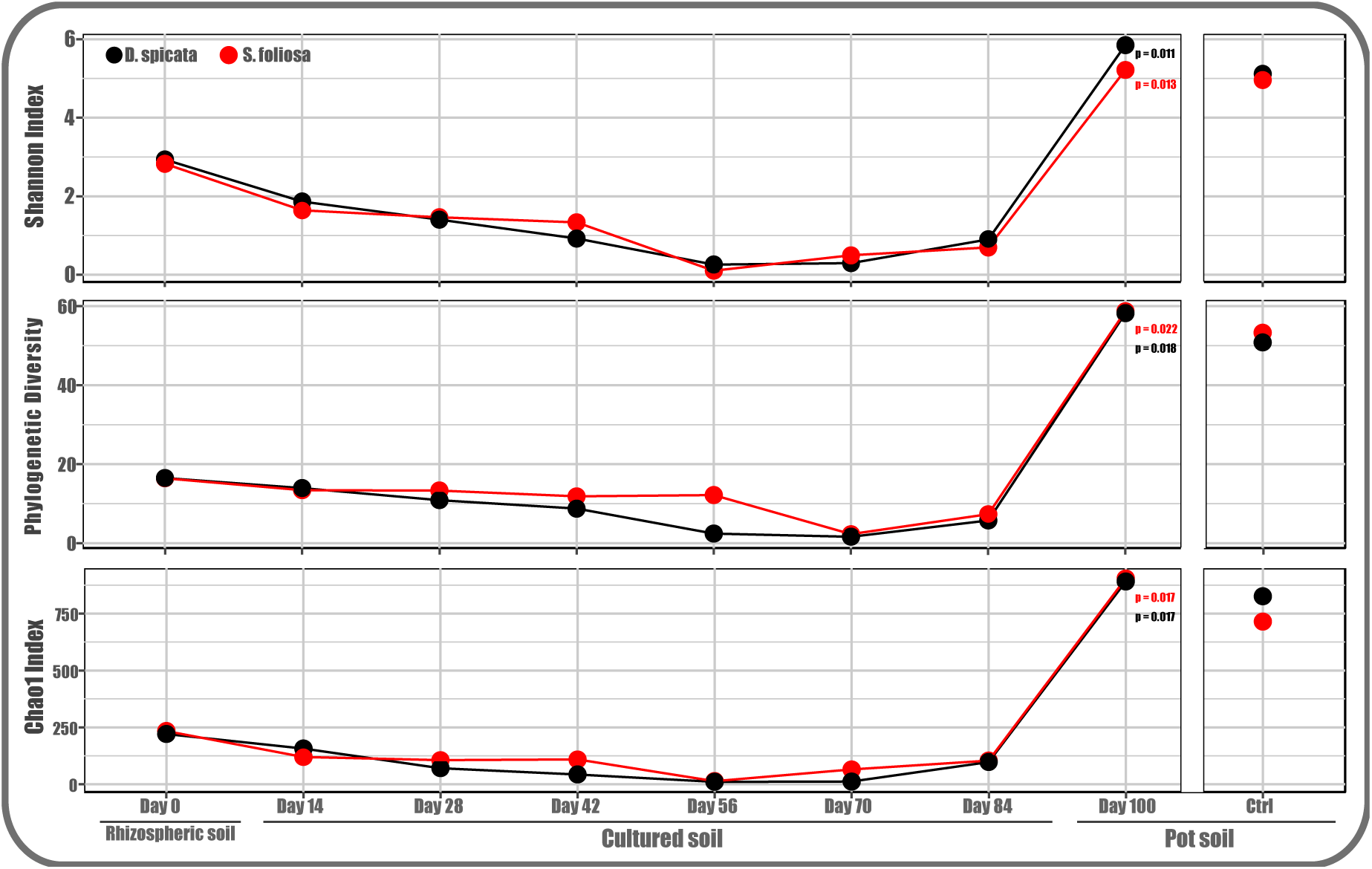
Alpha diversity monitoring through time and experimental stages. Faith’s phylogenetic diversity, Chao1 and Shannon indices were calculated for each evaluated point and their average values are shown in red for *S. foliosa* experiments and in black for *D. spicata*.

By analyzing the taxonomic composition throughout all stages of the experiment we were able to identify a total of 1,017 ASVs classified to the lowest taxonomic rank available (99.8% at Phylum, 99.02% at Class, 92.6% at Order, 82.9% at Family, 58.01% at Genus and 8.16% Species). The monitoring of taxonomic groups evidenced that over time some populations were selected and progressively enriched in the cultures, which is very clear within the Enterobacteriaceae family in both plants (Figure 3) that reached 95.2% and 98.4% of relative abundance by day 56 for *D. spicata* and *S. foliosa* cultures, respectively. This reflects the aforementioned decrease in diversity throughout time observed for the cultures of both plants. The Brevibacillaceae family is also enriched in the culture stage, reaching 18.6% and 43.5%, respectively for both plants. On the other hand, families such as Bacillaceae (25.4%) and Pseudomonadaceae (6.3%) which were the main representatives *S. foliosa* rhizospheric soils, were depleted throughout the culture period. Similarly, the Haloferacaceae (22.9%) and Porphyromonadaceae (26.7%) families disappear during *D. spicata* cultures.

**Figure 3.**
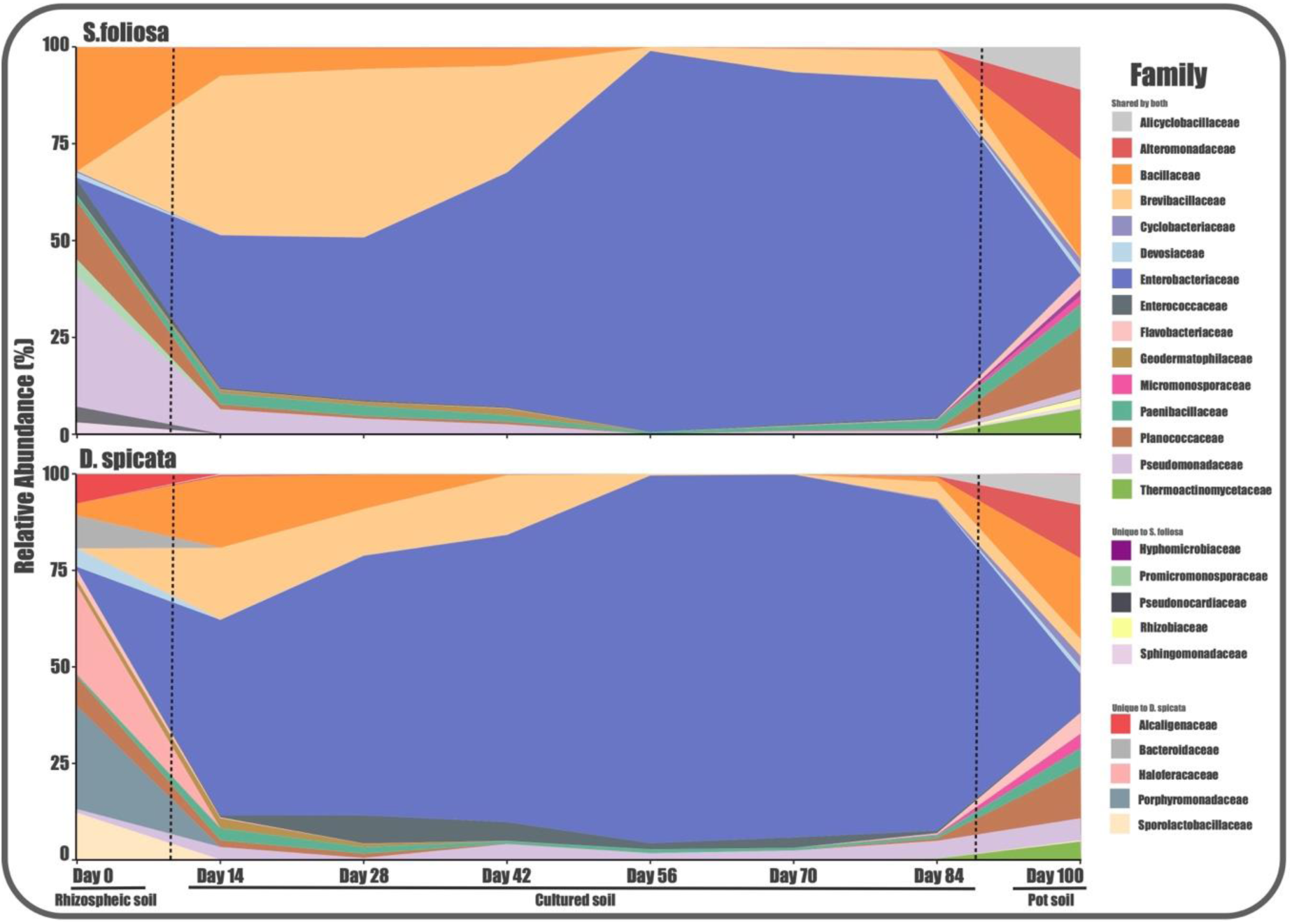
Taxonomic composition dynamics through time and experimental stages. Taxa were agglomerated to display the relative abundance of the top 20 family ranks for both plant species (code by colors).

Moreover, data shows that communities become more homogeneous during the culture stage regardless the soil sample origin, there were some differences among the detected families. The Hyphomicrobiaceae, Rhizobiaceae, Sphingomonadaceae, Promicromonosporaceae and Pseudonocardiaceae families were only detected in the *S. foliosa* rhizospheric soil, while the Porphyromonadaceae, Haloferacaceae, Sporolactobacillaceae, Alcaligenaceae and Bacteroidaceae families were unique for *D. spicata* rhizospheric soil. Also, the most abundant or dominant taxa belonged to the Pseudomonadaceae, Bacillaceae and Planococcaceae families for *S. foliosa* and the Porphyromonadaceae and Haloferacaceae families for *D. spicata*. All these results are reflected at the phylum rank with a significant presence and subsequent complete dominance by Firmicutes and Proteobacteria (Supplementary Figure S3).

The inoculation of lettuce plants with the 84 days cultures of both *S. foliosa* and *D. spicata* rhizospheric bacteria resulted in growth promotion of the plant, compared to those that were not inoculated with any type of culture or microorganisms. The supplemented ones were evidently larger and visually vigorous (Figure 4A). These differences were morphologically quantified considering the root length (Figure 4B), the leaf length (Figure 4C) and the accumulated dry matter percentage (Figure 4D). Notably, the lettuce plants that were inoculated with the cultures from *S. foliosa* rhizospheric soil showed a statistically significant increase in the three evaluated parameters. Alternatively, in those inoculated with the *D. spicata* cultures the promotion was in a less evident, nonetheless root length and dry matter content were significantly increased. Also, the lettuce plants inoculated with sterile culture medium did not show significant differences in any of the parameters regarding the control plants.

**Figure 4.**
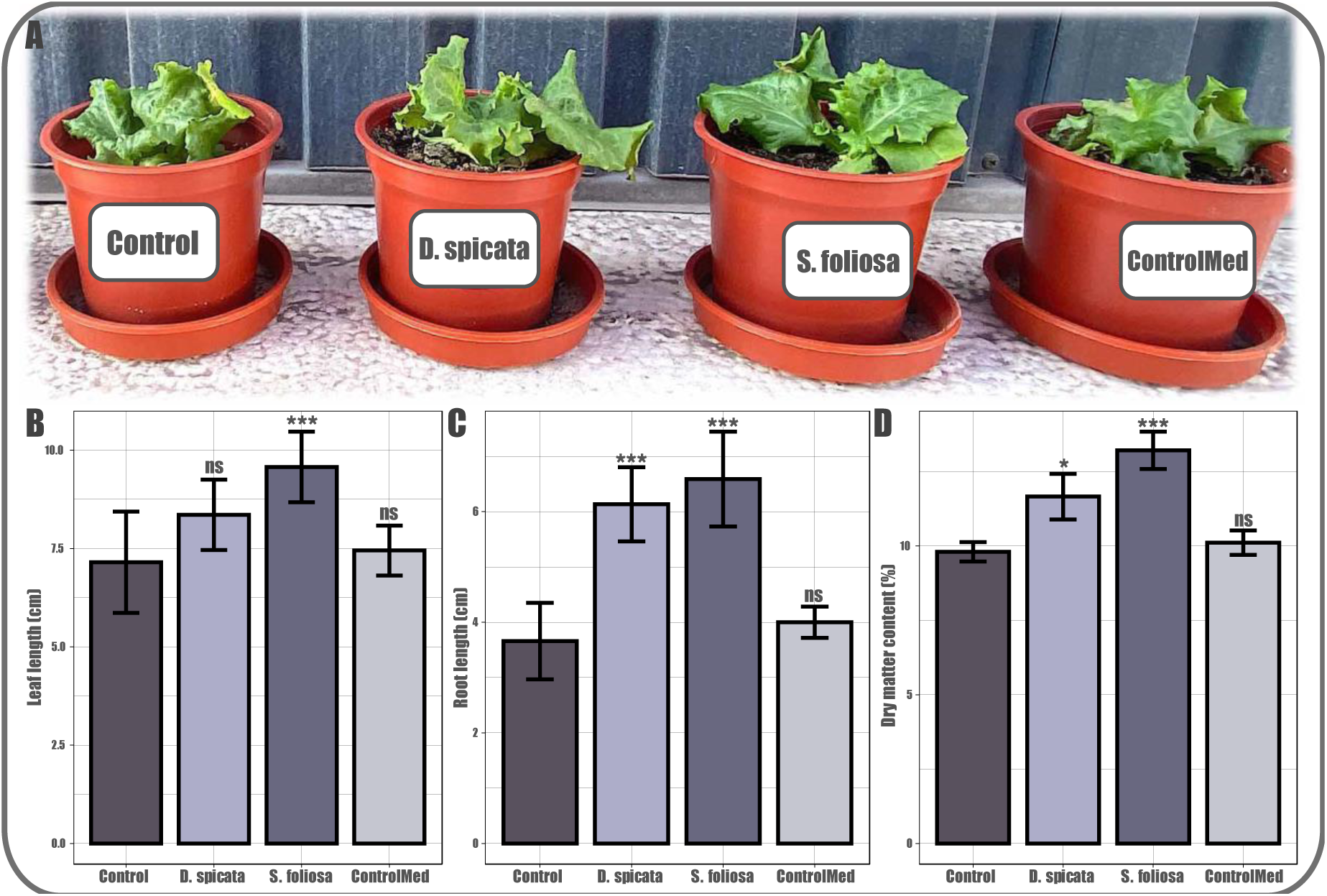
Effect of bacterial cultures inoculation on *L. sativa* growth. **A)** Day 100 lettuce plants picture in the four different treatments: Control (not supplemented); *D. spicata* (supplemented with the day 84 culture from the *D. spicata* rhizospheric soil); *S. foliosa* (supplemented with the day 84 culture from the *S. foliosa* rhizospheric soil) and ControlMed (supplemented with sterile culture medium). **B)** leaf length in cm; **C)** root length in cm and **D)** dry matter content in percentage. Presented data is an average of at least ten individuals. The symbols: * p ≤ 0.05; ** p ≤ 0.01; *** p ≤ 0.001, indicate statistical significance (regarding the control) according to ANOVA.

As we are aiming to determine which could be the key agents that promote the growth on the lettuce plants inoculated with the cultures, we set up to identify candidate taxa. When evaluating the communities composition agglomerated at genus rank, we observed mostly the enrichment of the *Klebsiella* and *Brevibacillus* genus as the culture progresses while the composition patterns change completely between the Yungay rhizospheric soils and the cultures with an evident decrease in diversity as well. We also noticed that the many of the detected families were represented by only one genus (Supplementary Figure S4). Moreover, other genera such as *Bacillus*, *Brevibacillus*, *Enterococcus,* and *Pseudomonas* were also detected in the cultures, although with a lower relative abundance. Also, comparing the communities composition between both plants rhizospheric soil, we can highlight great differences such as the dominance of *Porphyromonas* and *Haloterrigena* in *D. spicata*, as well as the great abundance of *Pseudomonas* and *Bacillus* in *S. foliosa*.

We identified 12 candidate ASVs that meet the selection criteria, of which 3 belong to the Proteobacteria phylum (ASV1: *Klebsiella* sp.; ASV4 and ASV17; which are two strains of *Pseudomonas* sp.) and 9 to the Firmicutes phylum (ASV7: *Paenisporosarcina* sp.; ASV20: *A. resinae*; ASV11: Enterococcus sp.; ASV40: *Bacillus* sp.; ASV27 and ASV74: two strains of *Paenibacillus* sp.; ASV80: a member of the Planococcaceae family; ASV14 and ASV162: two members of the Bacillaceae family) (Figure 5). Moreover, only 5 of the 12 ASVs were identified in the *D. spicata* experiments, while 11 of the 12 ASVs were identified in the *S. foliosa* experiments. Interestingly, one of the two Bacillaceae family members (ASV14) was exclusively detected on the *D. spicata* experiments, whereas ASV11, ASV17, ASV27, ASV40, ASV74, ASV80 and ASV162 were exclusively detected on the *S. foliosa* experiments. Additionally, 4 of the 12 candidate ASV are shared between both plants (ASV1, ASV4, ASV7 and ASV20). Interestingly, *Pseudomonas* sp. (ASV4) and *Bacillus* sp. (ASV40) which represented an important proportion in Yungay rhizospheric soils communities, decreased their abundance greatly throughout the culture stages. On the contrary, *Klebsiella* sp. (ASV1) who represent less than 1% of relative abundance in the Yungay rhizospheric soils communities, was enriched throughout the cultures until reaching virtually total dominance (98.37%).

**Figure 5.**
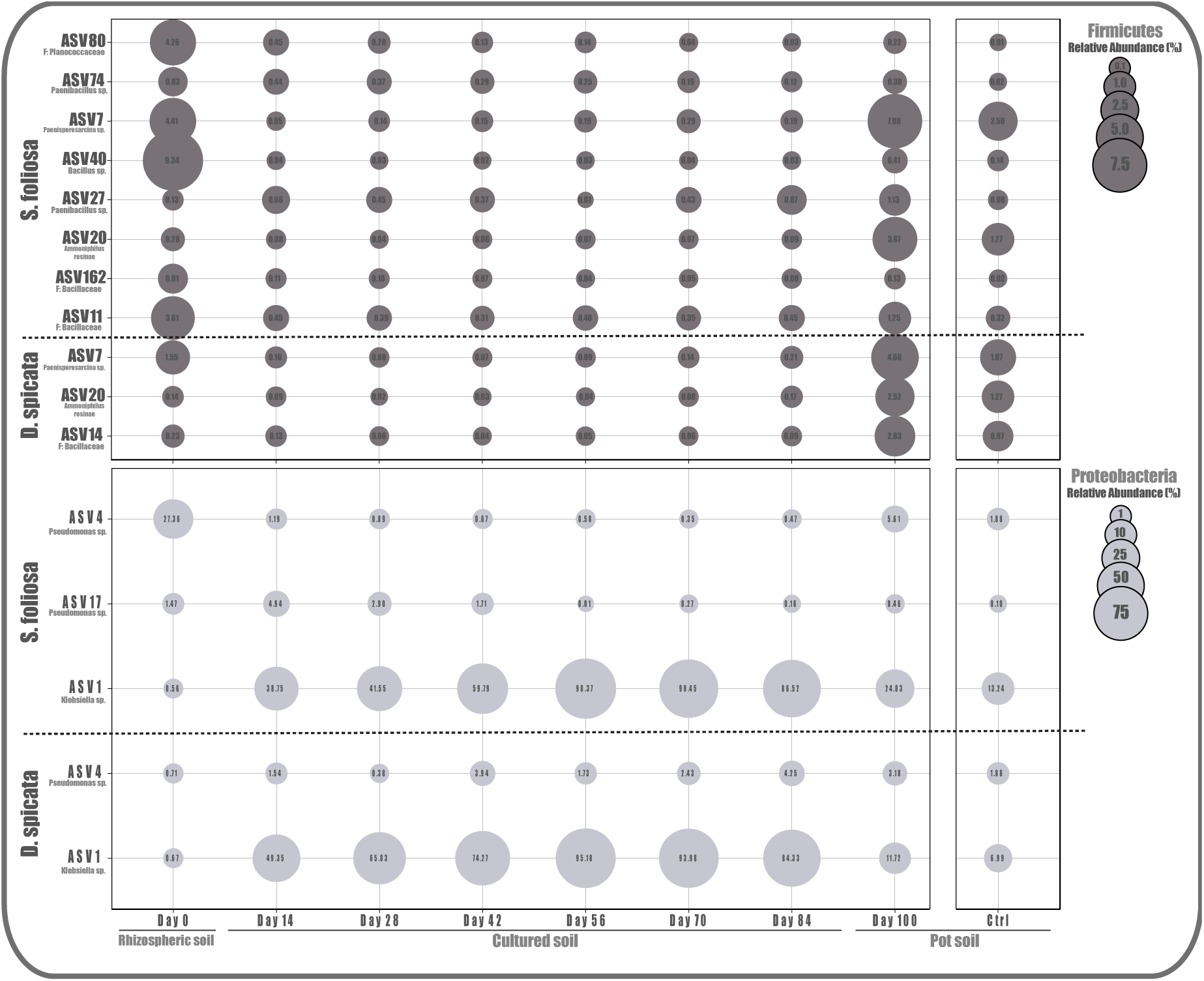
Potential key taxa identification. Taxa that met the established criteria are shown in the bubble plot, where each circle represents the relative abundance of each taxon (best available rank classification) at the corresponding experimental stage. Legends are color coded by phylum with the corresponding scale.

The co-occurrence network of the Lettuce pot soil communities was composed of 46 nodes (ASVs with at least one significant correlation) and 39 edges arranged in 9 modules (Figure 6). The modules are dominated by Firmicutes (17/46) and Proteobacteria (12/46), agreeing with all previous findings. Also, the nodes with the highest degree of interaction belong to ASVs of Firmicutes and Actinobacteriota, despite this last one being least abundant in the communities. Moreover, 5 of the 12 identified candidates are part of the network structure (ASV1: *Klebsiella* sp., ASV4: *Pseudomonas* sp., ASV7: *Paenisporosarcina* sp., ASV11: Bacialiaceae and ASV20: *A. resinae*), which are mostly Firmicutes and belongs to four of the nine network modules. Particularly, ASV1 (*Klebsiella* sp.) it is the central part of a module with four nodes, which may suggest an important structural role. Among its connections is ASV11 (*Enterococcus* sp.) which is another candidate and ASV2 (*Brevibacillus* sp.) which was one of the most abundant genera in culture stages. Finally, for these pot soils we identified 7 keystone species (ASV39: *Longispora* sp., ASV43: *Novibacillus thermophilus*, ASV53: *Ammoniphilus* sp., ASV55: *Planifilum* sp., ASV59: *Streptomyces* sp., ASV119: *Chryseolinea* sp. and ASV148: Gemmatimonadota) mostly belonging to Firmicutes and Actinobacteria phyla, which might be playing an important role in the community function, structure and stability.

**Figure 6.**
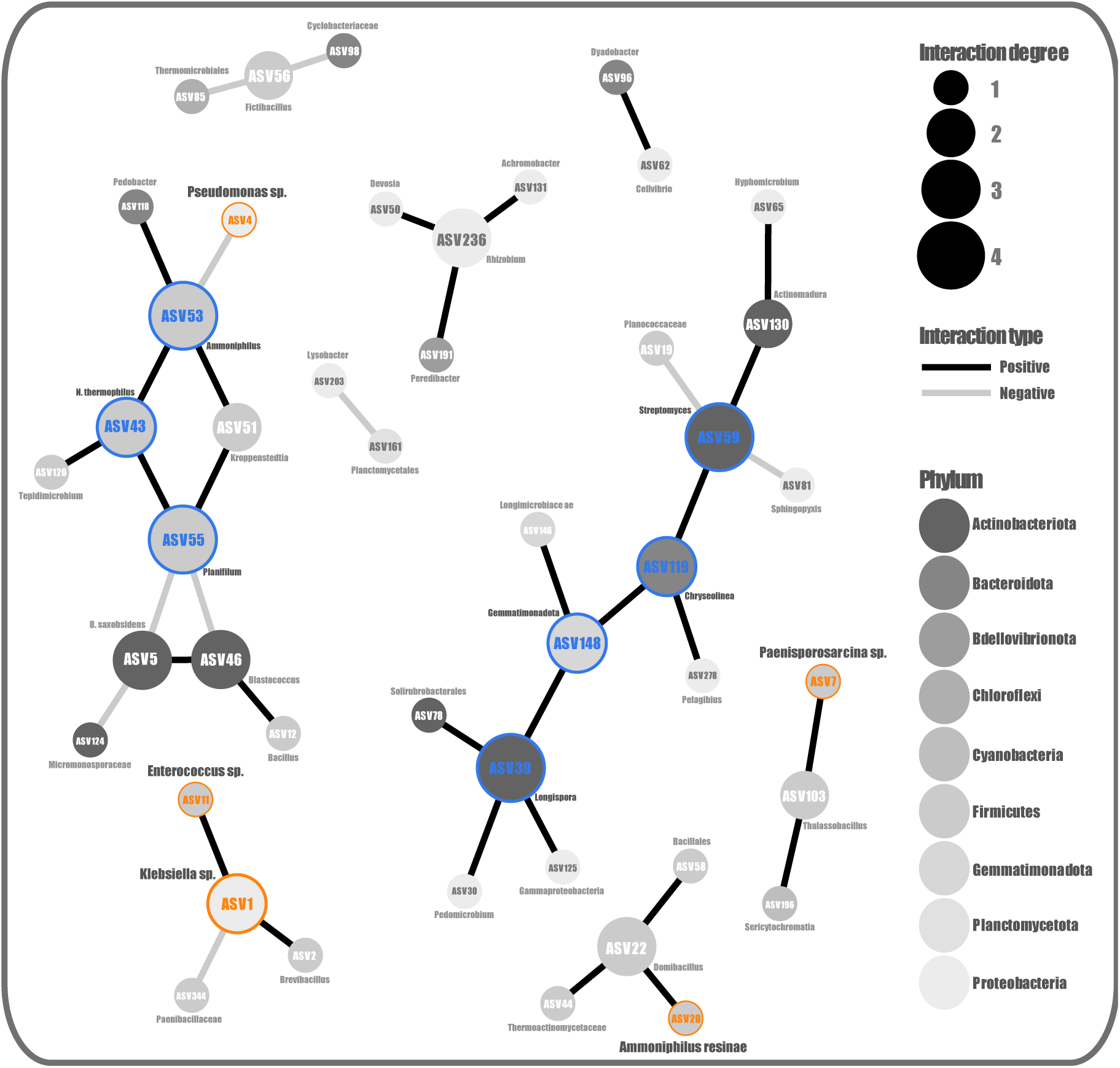
Co-occurrence networks of the *L. sativa* pot soil bacterial communities. The size of each node (representing ASVs) is proportional to the number of different interactions (degrees), the edges (significant connection between nodes) color represents the interaction type, the node color indicates the taxonomic affiliation at phylum level and node labels are at the lowest available taxonomic classification. The node border color denotes candidate species in orange and the identified keystone taxa in blue.

## DISCUSSION

In this work we were able to improve the growth of an agricultural relevant crop using standard and affordable microbiology methods to grow a bacteria consortium from the rhizosphere of plants thriving under the extreme abiotic conditions of the Atacama Desert hyper-arid core. This is the first report were the culturable fraction of a rhizospheric soil community from this extreme environment was monitored, describing it dynamics throughout time (from a taxonomical composition point of view). The microbial cultures were maintained over six subcultures for a total period of 84 days, and as expected, the community diversity decreased. The composition was enriched with the best adapted taxa to laboratory-controlled conditions, a critical aspect to consider for the development of biotechnologically applied tools, as the bacteria that are difficult to grow are not such attractive candidates for agricultural application.

The monitoring of the alpha diversity account for a selection process with a consequent and expected loss of diversity progressively. Although there were no significant differences between the two plant species, the decrease over time was significant for each of them. It is worth noting that *D. spicata* rhizobiomes composition exhibited less variability compared to *S. foliosa*, which accounts for a more stable or selected community. This may be related to the need for more specialized organisms capable of tolerating the high salinity and pH changes induced in the soil which are caused by the plant’s physiologic processes to eliminate salts and thus thrive under desertic conditions (Conticello et al., 2002; Pelliza et al., 2005). We also must consider that we are starting with low diversity levels in the rhizosphere soils, which is consequent and characteristic of Atacama Desert soils, particularly in the Yungay area (Neilson et al., 2017; Scola et al., 2018; Fuentes et al., 2022). On the other hand, the large increase in diversity observed in the pot soils (after the inoculations experiments) we believe is in mostly due to the lettuce plants own microbiome, which has been vastly described as having high richness and diversity (Žiarovská et al., 2022; Acin-Albiac et al., 2023), along with geographic and temporal variability (Yu et al., 2018). Also, the irrigation water and the environment could be input sources, even though the substrate was sterilized prior to the experiments, these were carried out in an open regime, since we tried to reproduce field conditions as much as possible.

The complete dominance or enrichment of *Klebsiella* genus and in a lesser extent the *Brevibacillus* genus reflects the selection process and account for what is seen at phylum (Proteobacteria and Firmicutes) and family (Enterobacteriaceae and Brevibacillaceae) ranks for both plants. Different species of the *Klebsiella* genus been recurrently detected in many types of soils and environmental samples (Ekwanzala et al., 2019), even in the soils of Yungay area (Thomas et al., 2016). Regarding the association of *Klebsiella* with plants, there are many reports of the great repertoire of beneficial traits provided: ACC deaminase activity, atmospheric nitrogen fixation, inorganic phosphate solubilization, great adhesion capacity (to the plant roots), production of indole acetic acid, siderophores, cellulase, protease and amylase enzymes as well as the promotion of saline, drought and oxidative stress tolerance; all which has been demonstrated experimentally, including irrigation tests with seawater (Rueda-Puente et al., 2003; Singh et al., 2015; Marasco et al., 2012; Acuña et al., 2019; Bakelli et al., 2022). Furthermore, the *Brevibacillus* genus has also been widely detected, isolated and cultured from the tissues and/or rhizospheric soils of plants inhabiting extreme environments such as the Siani, Kousséri and Atacama Desert and tested for plant growth promotion in crops such as tomatoes and corn. Among the beneficial capacities identified in these works are IAA and ammonia production, P-solubilization, ACC deaminase, extracellular enzymatic and antimicrobial activity (Soussi et al., 2016; Eke et al., 2019; ALKahtani et al., 2020; Astorga-Eló et al., 2021; Wang et al., 2022).

Both *Klebsiella* and *Brevibacillus* are easy to culture generalists, characterized by metabolic versatility and wide ranges of tolerance to abiotic factors, which could explain the exerted competitive exclusion during the culture on the R2A medium. The fact that these genera are easy to grow and work in laboratory-controlled conditions is probably the reason why there are so many studies on their abilities to promote plant growth (Rueda-Puente et al., 2003; ALKahtani et al., 2020). Also, all the previously mentioned characteristics and beneficial traits of these bacteria could account for the growth promotion evidenced our experiments with lettuce plants. Even though their abundance was not the majority in the pot soils, they clearly played an important role in the experiment’s outcome. The lettuce plants obtained beneficial capabilities from the cultures as evidenced in statistically significant increase in all parameters used to evaluate growth. Similar to the results obtained in previous works, where formulated/defined microbial consortia or isolated strains are used as biofertilizers to boost plant growth (Mondal et al., 2020; Santoyo et al., 2021; Fortt et al., 2022). Contrary to our approach, where a direct culture was used as a bioinoculant, which has not been reported before.

Even though the R2A medium was formulated for water samples it has been widely used to cultivate soil microorganisms, as it promotes low-growing heterotrophic bacteria (Chaudhary et al., 2019). It has been demonstrated that this medium capture much more diversity compared to many other widely used ones (Blickfeldt, Brain Heart Infusion, Frazier, Trypticase Soy, Lysogeny Broth, Nutrient and Yeast Extract) and it use has been promoted for the metabolomic profiling of soil bacteria (Dziurzynski et al., 2020; de Raad et al., 2021). Therefore, we believe that this culture medium and conditions promoted a competitive advantage for *Klebsiella* and *Brevibacillus*, since both are generalists, versatile and adaptable (Bakelli et al., 2022; Wang et al., 2022). This, added to the fact that they are easily isolated, cultured and to manage in the laboratory, make them ideal biofertilizer candidates.

Although taxonomic composition of the cultures ended up being equivalent, there were differences between the rhizosphere soils of both plants; *Pseudomonas*, *Bacillus* and *Paenisporosarcina* were the main genera of *S. foliosa*, while *Porphyromonas* and *Haloferax* dominate *D. spicata* rhizobiome, all of which were depleted by competitive exclusion during the culture stages. *Pseudomonas* has a long-lasting relation with plants and is recurrent in desertic environments, being vastly reported its capabilities to colonize plant surfaces and inside tissues, thus promoting plant growth by suppressing pathogens and synthetizing phytohormones (Preston, 2004; Gaete et al., 2022). Nonetheless, some species as *P. syringae* is a well-known plant pathogen (Xin et al., 2018). Moreover, *Bacillus* genus is well known for its versatility, stress tolerance and its ability to form very resistant spores, particularly its interactions with plants have also been widely studied, demonstrating its ability to colonize the roots through biofilm formation, stimulate growth, as a biocontrol agent and facilitating tolerance to abiotic stress (Hashem et al., 2019; Tsotetsi et al., 2022). This agrees with our findings due to the Yungay area conditions and reaffirms the relevance of these genera for agriculture improvement in the context of a climate change scenario. Also, bacteria from the *Paenisporosarcina* genus (previously classified as *Sporosarcina*) have been identified as Gram-positive, spore forming, generalist, coccobacillus and some can be psychrophilic (Reddy et al., 2013). This genus has been recurrently detected and cultured from the rhizosphere of different plants in environments such as the Bolivian Altiplano and the Luoyang province in China, presenting a wide range of tolerance to temperature, drought, limited carbon sources and even heavy metals (Han et al., 2011; Gomez-Montano et al., 2013). Interestingly, this genus has also been reported as the second most detected endophyte of *Atriplex* spp. in the Kalahari Desert and Jornada del Muerto; this plant belongs to the Chenopodioideae subfamily along with *S. foliosa* (Tahtamouni et al., 2016).

The archaea genus *Haloferax* characterized as a denitrifying halophile was also detected as a majority component of *D. spicata* Rhizobiome, this microorganism has also been associated with benefits for plants, particularly the production of phytohormones such as IAA (Indole Acetic Acid) that promote growth and siderophores which contribute to change the soil physiochemical properties and mitigate stress (Ma et a., 2016; Yadav et al., 2017; Selim et al., 2022). Additionally, we detected the genus *Porphyromonas* in high abundance, which considered a pathogen (Guilloux et al., 2021), although there are some reports of this genus in environmental samples (Acuña-Amador and Barloy-Hubler, 2020), we did not find any report that associates this anaerobic bacterium with plants or any beneficial capacity.

By tracking the taxa present in the rhizosphere soils of Yungay, that persisted during culture and were detected in the pot soils after the biofertilization experiment, we identified microorganisms belonging to the genera *Klebsiella, Pseudomonas, Paenisporosarcina, Ammoniphilus resine, Enterococcus, Bacillus* and *Paenibacillus* and the families Planococcaceae and Bacillaceae. As well as those mentioned above, these organisms have been described previously as having beneficial capacities for the plants with which they interact. Particularly, *Enterococcus* produces different phytohormones and can promote tolerance to salt stress (Lee et al., 2015; Panwar et al., 2016). On the other hand, *Paenibacillus* has the ability to fix nitrogen and also inhibit phytopathogen nematodes (Khan et al., 2008; Liu et al., 2019). On the other hand, the *Ammophilus* genus is very interesting because it is an oxalotrophic bacteria that can secrete organic matter hydrolases to accelerate substance degradation and promote nutrient recycling (Wang et al., 2022). Although there are no reports that associate *Ammoniphilus* with plant growth promotion if it has been detected in rhizosphere soils (Yadav and Saxena, 2018; Abuauf et al., 2022). Interestingly, two members of this genus are part of the co-occurring community, being one of them also identified as a keystone species. Moreover, candidates *Klebsiella, Pseudomonas, Paenisporosarcina,* a *Bacialiaceae* member were also part of the co-occurring community, which agrees with the findings and imply an important role for these taxa that worth continue to investigate.

The other identified keystone species (*Longispora* sp., *Novibacillus thermophilus*, *Planifilum* sp., *Streptomyces* sp., *Chryseolinea* sp. and a Gemmatimonadota member) may also be targets for more mechanistic research, but we must consider that the origin of some of these could be the lettuce plants own microbiome (Cipriano et al., 2016). Finally, as the growth promotion effect was greater in the lettuce plants inoculated culture generated from the *S. foliosa* rhizospheric soil we would like to point out that this culture 9 of the 11 identified candidates were present, while in *D. spicata* only 5 of the 12 were detected. Furthermore, also in the *S. foliosa* cultures there were fewer *Pseudomonas* was less abundant and *Paenibacillus* was more abundant, which can also give us clues to discover which organisms may have more influence in the observed results.

## CONCLUSION

The relevance of this investigation is the direct use of cultures generated from the rhizospheric soil of plants thriving under the harsh conditions of the Atacama Desert Hyperarid Core. Clearly the culturable fraction of these rhizobiomes is able to transfer key traits to the crop to improve its growth and yield, which is very appealing as the use of a small group of microorganisms that are easily grown in laboratory-controlled conditions can have a significant and positive effect over an economically important crop. It is important to mention that all these experiments were carried out in the city of Antofagasta where the desertic climatic is maintained therefore the plants were subject to these variables even though they were maintained with constant watering and in the shade. Also, we propose as the next step, to isolate *Klebsiella* strains and test their plant growth promoting effect a as monoculture due to their possible role on the obtained results. Finally, we want to highlight the novelty of the work and relevance of the results obtained, this being the first report where the taxonomic composition of soil cultures is monitored over time and subcultures and for the evaluation of bacterial consortia from the Atacama Desert native plants as good biofertilizers.

## AUTHOR CONTRIBUTIONS

JC-S, JF and FR conceived and designed the study. JC-S, JF, GD, PA, MS and FR performed the field work. JC-S, PA, MS, GD, JF and MA processed the samples and performed the experimental procedures. JC-S carried out the bioinformatics analyses. FR, CS and AS contributed with reagents, materials, and analysis tools. JC-S, CP-E, AS and FR interpreted the results and wrote the first manuscript draft. All authors read and approved the final manuscript.

## ACKNOWLEDGEMENTS AND FUNDING

This research was sponsored by ANID (Agencia Nacional de Investigación y Desarrollo de Chile) grants. 2021 Postdoctoral FONDECYT 3210156, 2022 Regular FONDECYT 1220902 and 2022 Estrategia en Sequía FSEQ210029 to JC-S and FR. 2021 Regular FONDECYT 1210633 and 2023 Anillo ATE220007 to CPS. 2023 Postdoctoral FONDECYT 3230189 to CP-E. The funders had no role in study design, data collection and analysis, decision to publish, or preparation of the manuscript.

